# FE-DOE: Finite element informed design of experiments to optimise bioinspired melt electrowritten (MEW) polymeric heart valves

**DOI:** 10.1101/2025.11.01.684363

**Authors:** Celia Hughes, Robert D. Johnston, Desmond McCarthy, Ewa Klusak, Emily Growney, Evelyn Campbell, Caitríona Lally

**Author notes:** Corresponding author: Caitríona Lally; Parsons Building, Trinity College Dublin, College Green, Dublin 2, D02 PN40.

## Abstract

Aortic stenosis is predominantly treated through transcatheter bioprosthetic heart valve implantation. The materials used in these devices, however, suffer from premature failure. Polymer heart valves have the potential to improve current commercial devices, offering materials with extended durability and customisation through fibre-reinforcement. Due to the wide range of available materials and structures, there is a need for a methodical approach to the design and optimisation of novel, bioinspired polymeric leaflets. This work presents a framework using FE and DOE tools to enable the creation of optimised bioinspired, 3D-printed, fibre-reinforced polymer leaflets using MEW. Here, FE models are created to represent MEW fibre-reinforced polymer leaflets for application in a transcatheter aortic heart valve. The behaviour of this valve under physiological loading conditions is modelled to predict valve performance and leaflet material response. These models were first used to investigate the impact of fibre orientation on valve performance and leaflet response, showing the benefit of using a bioinspired fibre reinforcement structure. Using DOE, the structural combination of MEW fibre-reinforcement and elastomeric matrix was optimised based on valve performance and leaflet stress and strain. Overall, the framework offers an efficient and versatile methodology for optimising fibre-reinforced polymer leaflets by utilising an in-silico approach to remove the need for manufacturing and testing of these devices.

## Introduction

Aortic stenosis is a prevalent disease of the aortic valve, affecting 4% of people over the age of 70 in the US, and leads to a 50% chance of mortality within two years if left untreated (Ambrosy et al. 2023; Généreux et al. 2023). Where surgical methods are not indicated, most treatment is achieved through minimally-invasive means using transcatheter aortic valve implantation (TAVI) (Carroll et al. 2020). In this case, the valves implanted contain leaflets made from biological materials, either porcine or bovine pericardium, and can suffer from premature failure due to material insufficiency and calcification (Dvir et al. 2018; Schoen and Levy 2020). Pericardium is a stiff, collagenous tissue that could potentially exhibit improved durability in TAVI devices if the fibres within the pericardial leaflets are aligned to bear the load (Guerin et al. 2025; Whelan et al. 2021). However, robust fibre screening methods are not currently utilised or required for the manufacture of these devices (Whelan et al. 2019), and it would be challenging to source tissue with an idealised fibre structure for leaflet load bearing.

Polymer heart valves offer the potential of a durable and biocompatible alternative to current commercially available devices. There are a wide variety of polymer heart valves currently in development which have taken unique approaches to the creation of these next-generation devices. These have primarily focused on developing novel polymer blends, such as the Tria valve (Dandeniyage et al. 2017; Jenney et al. 2021), PolyNova valve (Kovarovic et al. 2023; Rotman et al. 2020), PoliValve (Stasiak et al. 2020), a HA-LLDPE valve (Khair et al. 2025), and Life Polymer (Scherman et al. 2019). Notably, the Tria valve has recently become the first ever commercially available polymer valve approved for use, gaining approval in India for use in the mitral position (Foldax.com 2025).

Some polymer valve development has also been focused on the integration of a customisable fibre-reinforcement structure. This can be controlled using novel polymers, such as the copolymer blocks used in the PoliValve that can be arranged to give a mechanical orientation (Stasiak et al. 2020). Fibres can also be applied by leaflet additions such as the 3D-printed valve developed by Coulter, et al. which has bioinspired fibres overlaid on a polymer base (Coulter et al. 2019; Zeugin et al. 2024). Commissural and free edge fibres are also added in new generations of a HA-LLDPE valve (Khair et al. 2025). Additionally, embedded leaflet reinforcements have been explored with NiTi fibres to support leaflet stresses in the radial direction (Chen et al. 2023) and integration of a nylon fibre mesh (Choi et al. 2024). Many of these studies have shown reductions in leaflet strains and improved hydrodynamic performance with these fibrous inclusions (Chen et al. 2023; Coulter et al. 2019; Giaretta et al. 2024; Khair et al. 2025).

In addition, there has been considerable development in bioinspired, tissue engineered valve leaflets (Boehm et al. 2023; De Jesus Morales et al. 2024; Saidy et al. 2019; Snyder and Jana 2023). While these devices still require many years of development before they can be made commercially available, they have shown potential for adopting manufacturing methods such as melt electrowriting (MEW) for the creation of a bioinspired fibre reinforcement structure (Saidy et al. 2019). MEW is a highly controllable extrusion-based additive manufacturing technique that can produce structures with fibres at a biologically-relevant scale (Brown, Dalton, and Hutmacher 2016).

Previous work in our lab has studied the mechanical response and collagen microstructure of native porcine aortic valve leaflets (Hughes et al. 2025). Using this as a baseline, MEW can be used to manufacture a bioinspired fibre-reinforcement structure that can be embedded in a compliant elastomeric matrix to mimic the structure and mechanical response of aortic valve leaflets.

Finite element (FE) modelling is a powerful and versatile tool which can be used to aid the design and development of polymer valves. It has been used primarily to improve already-existing designs by modifying leaflet thickness, shape, or adding simple reinforcement structures (Chen et al. 2023; Coulter et al. 2019; Khair et al. 2025; Kovarovic et al. 2022; Masheane, Preez, and Combrinck 2024), with the potential to transform the design, development, and assessment process of these devices. By leveraging FE modelling as a tool, we can assess novel material combinations and structures in silico to guide future manufacture and bench testing. Implementation of these tools would dramatically decrease development time for these devices by reducing manufacturing and assessment time of potential material combinations.

Design of experiments (DOE) is another valuable tool that can be leveraged in the development of novel valve leaflets. With a seemingly endless range of materials and structural features to choose from when designing composite, bioinspired leaflets, DOE can aid the process by providing a systematic approach to efficiently test combinations of features to enable sound statistical analysis of each feature’s impact (Durakovic 2017). DOE has been used alongside FE modelling to improve manufacturing workflows and optimisation for applications such as rotary endodontic instruments (Di Russo et al. 2025), circular saw tooth geometry (Schreiner et al. 2023), and steel bridge stiffeners for bridge assembly (Navarro-Manso et al. 2015). To the authors’ knowledge, while the combination of FE modelling and DOE can provide an efficient design workflow, it has not yet been used in polymeric valve development.

The objective of this work was to develop an integrated FE modelling and DOE approach for the design of novel, bioinspired, fibre-reinforced polymeric valve leaflets. Based on structures that can be manufactured using MEW, it aims to determine the best combination of printed-fibre structural features embedded in a compliant elastomeric material. Additionally, it aims to verify the benefit of a fibre-reinforced structure with bioinspired local orientations over other reinforcement types. Ultimately, this design framework should be usable with any material combination to determine the best combination of structural features for prosthetic aortic valve leaflets.

## Methods

In order to simulate a fibre-reinforced leaflet, first a model with discrete fibre inclusions was developed to obtain the mechanical response of a MEW/ elastomer composite. This response was used to calibrate a hyperelastic, fibre-reinforced constitutive model to describe the material response and fibre orientations in a trileaflet valve model. This methodology was first used to verify the benefit of bioinspired fibre orientations, and then it was used alongside a DOE approach to determine the best MEW and elastomer structural combination.

In this work, the materials simulated were Sylgard-184 (Dow Inc., MI, USA) polydimethylsiloxane (PDMS) at a 16:1 base-to-curing volume ratio embedded with a polyether ether ketone (PEEK)-like fibre structure.

### Discrete fibre model

A dogbone-shaped model with discrete fibre inclusions was created in Abaqus (SIMULIA, Dassault Systèmes, 2022) to simulate the mechanical response of a MEW-embedded elastomer. To create this, a dogbone model and a MEW mesh were created separately. The dogbone shape had an 18 mm gauge length, 4.5 mm width, and variable thickness. A MEW mesh was created in SolidWorks (SolidWorks Corp., Dassault Systèmes, 2023) with an assumed circular fibre diameter of 20 µm. The mesh cross section was created using a slot shape, with a thickness of 20 µm and a height of 20 µm x total layer number. Layer number and fibre spacing were variable. All of the layers were modelled as a solid part, see Figure 1.

**Figure 1.**
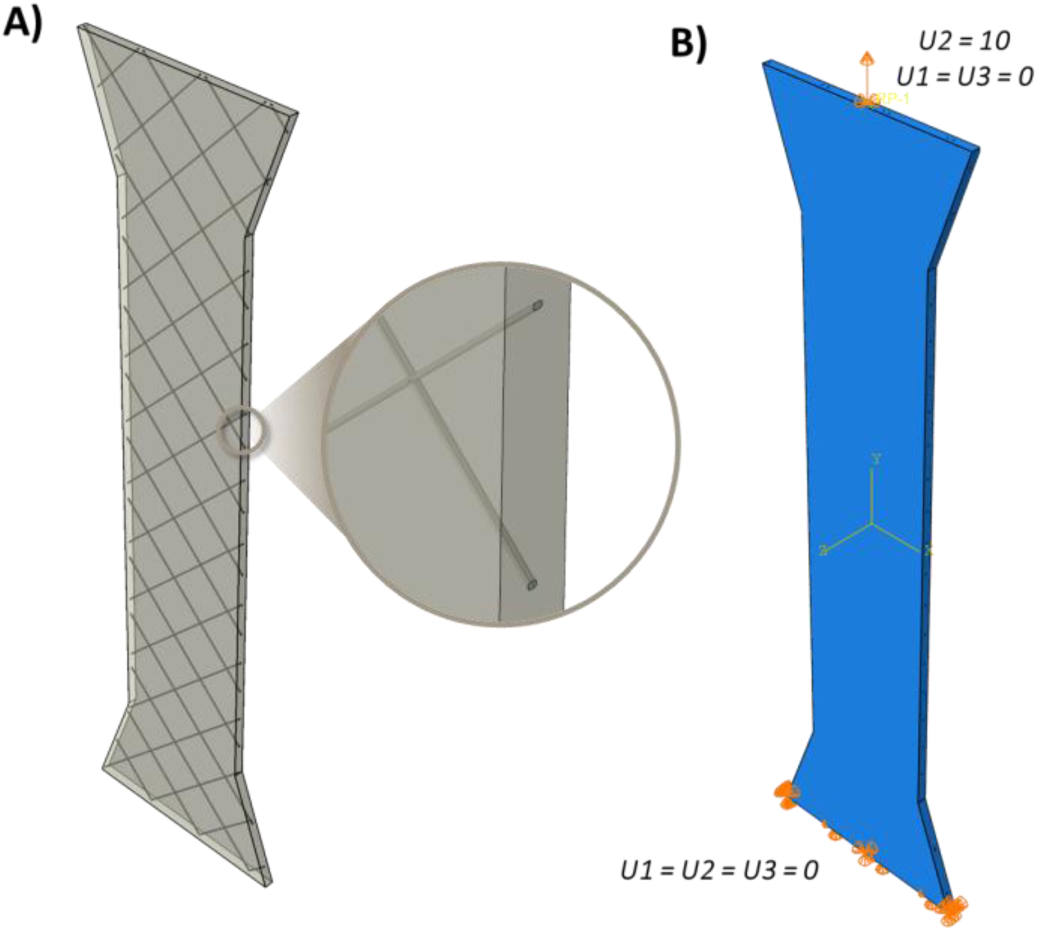
Uniaxial dogbone model with discrete fibre inclusions showing (A) fibres embedded in matrix with different material assignments, and (B) model boundary conditions.

In Abaqus CAE, the two parts were merged together with the boundaries retained. Each component was assigned unique material properties, and it was meshed as one solid part. For mesh generation, the part was seeded with an element size of 0.0625 mm for the fibres and 0.2 mm on the outside edges. It was meshed with 4-node linear tetrahedron elements. The matrix component, PDMS, was modelled as a first order Ogden material with a µ1= 0.204 MPa, α1= 2.504, and D=0; material parameters were calibrated from previous in-house uniaxial tensile testing. A mesh sensitivity analysis and material calibration curves can be found in the supplementary material. The fibres were modelled as a linear elastic material based on PEEK with a Young’s modulus of 2,906 MPa and a Poisson’s ratio of 0.45 (Shrestha et al. 2016). To simulate a uniaxial test, the bottom surface of the dogbone was fixed (U1=U2=U3=0) and the top surface was constrained using kinematic coupling to a reference point; all degrees of freedom were constrained (U1, U2, U3, UR1, UR2, UR3). A displacement boundary condition of 10 mm was applied to the reference point to deform the model (U2=10; U1=U3=0).

The true stress/strain response of this material was calculated using force/displacement output from the reference point, and used to calibrate the in-built Gasser-Ogden-Holzapfel (GOH) model in Abaqus using MCalibrate (PolymerFEM, MA, USA) (Gasser, Ogden, and Holzapfel 2006).

### Trileaflet valve model

A trileaflet valve model inspired by the ACURATE neo2 Aortic Valve™ (Boston Scientific, Marlbourough, Minnesota, US) was developed to model the behaviour of MEW-embedded polymer leaflets in a simulated valve environment.

#### Geometry, mesh, material, and fibre orientations

For the valve model, the leaflet geometry was designed based on the large size ACURATE neo2 Aortic Valve™. The geometry was drawn in Abaqus CAE and partitioned into 56 discrete rectangular regions to later assign local fibre orientations. After partitioning, the leaflet was exported as an STL file and imported into ANSA Beta (Beta CAE Systems, MI, USA) for uniform meshing into linear, full integration brick elements (C3D8), at which point it was imported back into Abaqus as an INP file. After performing a mesh convergence study (see supplementary), the optimum number of elements was selected at 22,715. Each of the 56 regions was assigned a unique local fibre orientation for two families of fibres, see Figure 2. These orientations were informed based on the measurements obtained by Hughes, et al. 2025: orientations for the belly region were assigned to a partition in the same region on the leaflet model and the commissure orientations were assigned to a partition in the corresponding region (Hughes et al. 2025). Fibre orientations were extrapolated based on these for the rest of the leaflet, ensuring no family orientation differed by more than 5° from its neighbouring regions. Material behaviour in the leaflet was described by the in-built GOH model, with parameters calibrated from the discrete fibre uniaxial model.

**Figure 2.**
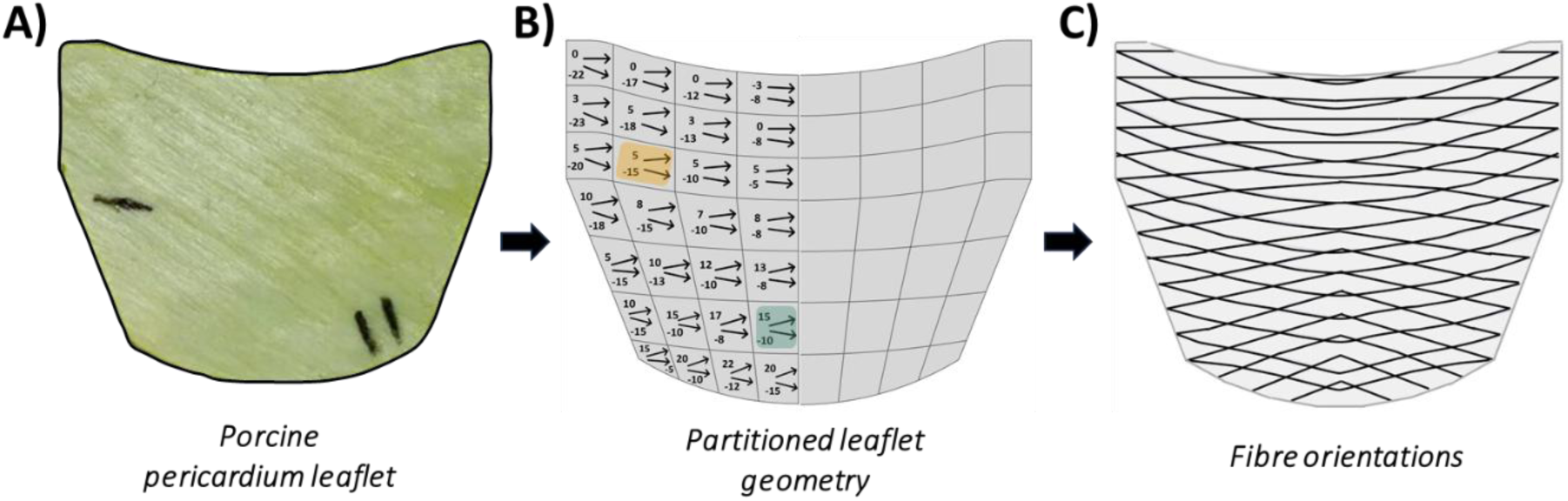
Porcine pericardium leaflet from ACURATE neo2 aortic valve compared to (B) leaflet model including partitions and local fibre orientations, and (C) diagram of fibre orientations across leaflet. Orange and blue regions in (B) correspond to fibre orientations informed from imaging of native leaflets’ commissures and belly regions (Hughes et al. 2025).

To simulate the frame and enable accurate leaflet “suturing”, a surface part was created representing the geometry of the frame, see Figure 3A. This was meshed with 2,608 SFM3D4R elements. To assemble the whole model, three leaflets and three frame surfaces were brought in at 120° to each other to form a symmetrical valve. Local coordinate systems were defined for each leaflet-frame surface pairing in order to aid boundary condition definition.

**Figure 3.**
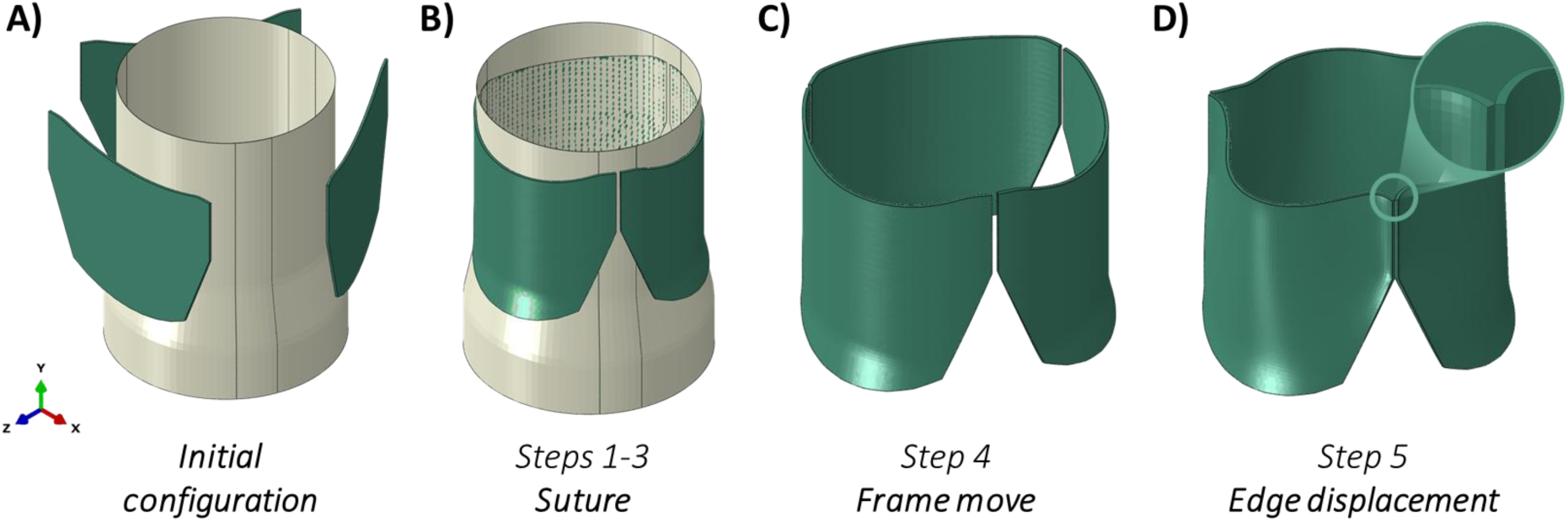
Valve model setup steps before pressure application to simulate sutured leaflet.

#### Boundary conditions and steps

The valve was modelled using ABAQUS/Explicit solver. There were 8 steps total in the model comprised of four steps to suture the leaflet, one to move the frame out of contact, and three pressure steps, see Figure 3.

Steps 1, 2, and 3 moved the leaflet into contact with the frame, moved the edges close to the frame, and then pressurised the leaflet against the frame, respectively. Throughout these three steps the frame was fixed in place (U1=U2=U3=0). In Step 1, the entire leaflet moved towards the frame (U3= −1.3 mm) and was constrained in the other two directions (U1=U2=0). In Step 2, the sides of the leaflet moved towards the frame (U3=-2 mm, U2=0) and the entire midline of the leaflet was fixed in place to stabilise the part (U1=U2=U3=0). For Step 3, a pressure was applied to the leaflet outer surface until it was fully in contact with the frame (p=0.005 MPa). To prevent vertical or side-to-side translation of the leaflet, a portion of the midline was constrained (U1=U2=0), see Figure 3B.

At this point, the frame was no longer needed and was moved away from the leaflet (U3=-20 mm), see Figure 3C.

To complete suturing, the inner edges of the leaflets should come into contact, as they would in a fully assembled valve, see Figure 3D. To simulate this, a displacement of total leaflet thickness in the X-direction was applied on these edges. Finally, the leaflets were in their sutured configuration and the pressure was applied. For a detailed view of boundary conditions applied throughout the model steps, see supplementary material.

Pressure was applied as a dynamic load to both the aortic and ventricular leaflet surfaces, as shown in Figure 4A, to simulate the opening and closing of the valve during a single cardiac cycle (Sacks, David Merryman, and Schmidt 2009). These curves, provided by Boston Scientific, were extracted from pulse duplicator testing on size large ACURATE neo2 devices that were tested using a ViVitro Pulse Duplicator system (ViVitro Labs Inc., BC, Canada) under hypertensive pressure (peak trans-aortic pressure of 140 mmHg) and nominal flow conditions (70 bpm, 35% systolic duration, and 5 l/min), as outlined in ISO 5840. One full cycle, lasting 0.857 seconds, was applied across a single step. To minimise the effect of the initial geometry and any inertial effects from the previous step, the pressure conditions was repeated three times across Steps 6, 7, and 8, and results were taken from the final step.

**Figure 4.**
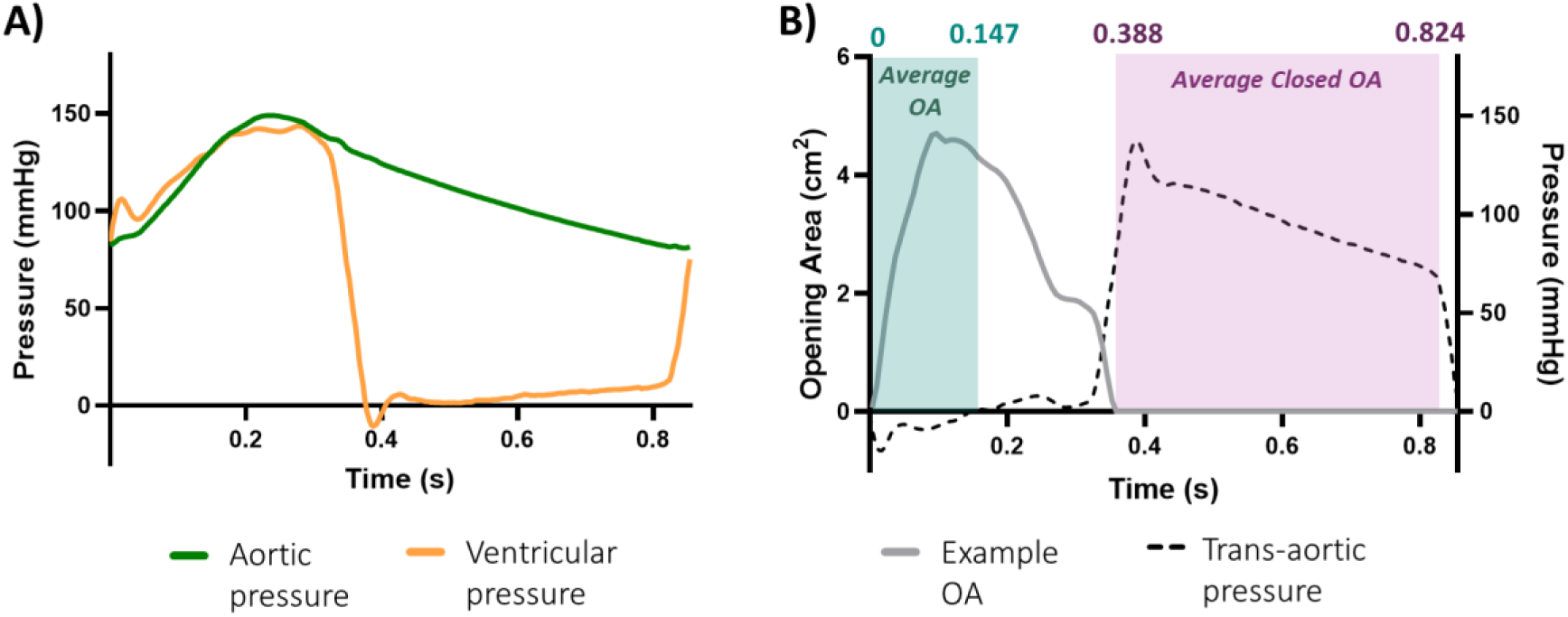
(A) Pressure applied to aortic and ventricular surfaces across a step, and (B) diagram showing resultant aortic pressure alongside an example OA curve detailing the regions taken for average OA and average closed OA calculations.

Contact for the leaflet-stent and leaflet-leaflet interactions was defined using the general contact algorithm. The same interaction property was assigned to all pairs: tangential behaviour was formulated using the penalty algorithm with a friction coefficient of 0.1, and normal behaviour was defined as “hard” contact with the default constraint enforcement method. Separation was allowed after contact.

#### Valve opening area extraction

A custom python script was created to extract images of the top view of the valve during its final cycle, amounting to 128 images each representing a time step of 0.0067 s. These were analysed in a custom MATLAB (MathWorks Inc. (2024), MA, USA) script that used edge detection to calculate the opening area of the valve, see Figure 5. Using the known dimensions of the valve, this was converted from pixels^2^ to cm^2^. With these outputs, a variety of parameters were extracted, see Figure 4B. First, maximum opening area (OA) across the entire cycle was identified. However, since this is just one measurement taken from a period with varying areas, a secondary average OA was calculated to serve as a surrogate for effective orifice area (EOA). EOA is a clinical metric of valve opening performance derived from flow measures, and for valves of this size it is required to be at least 1.7 cm^2^ (ISO 5840-3). Average OA was taken as the mean area during the time when ventricular pressure is higher than aortic pressure, which corresponded to 0-0.147 s. This aligns with methods employed by our commercial partners, where EOA during pulse duplicator testing is taken based on the cardiac output across this region. This measurement was validated by modelling porcine pericardium leaflets and comparing the average OA measurement to expected EOA values and these values were found to align well (see supplementary). Also extracted was a measure of closure, which was taken as an average OA during the end of the cycle (0.388 - 0.824 s). Finally, as a measure of valve “efficiency”, the time from the start of the cycle to maximum opening area was identified.

**Figure 5.**
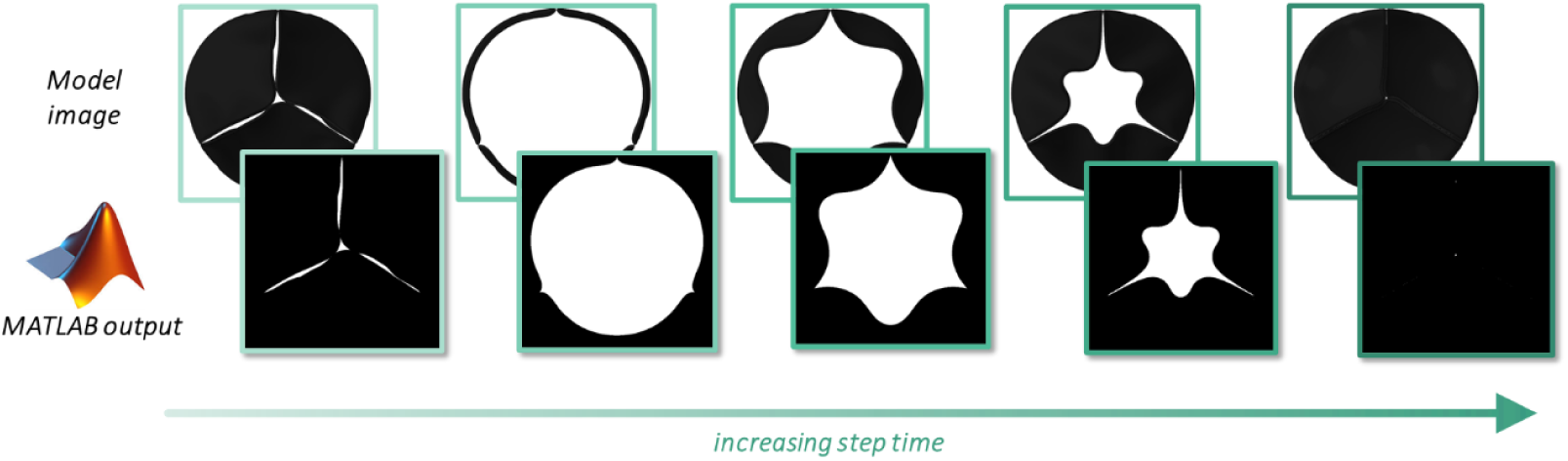
Diagram comparing image of valve from the top view extracted from Abaqus to the output from custom MATLAB script to determine valve opening area.

### Fibre orientation verification

In order to determine the impact of using bioinspired fibre orientations, the leaflet material response and valve performance were compared for different fibre reinforcement structures. For this, five models with different fibre configurations were investigated: bioinspired fibre orientations, all circumferential, all radial, isotropically dispersed, and no fibres. The material models in all of the fibre reinforced structures were calibrated based on a 2-layered MEW structure with fibre spacing of 2 mm embedded in 0.3 mm of PDMS. The calibrated GOH parameters for this were: C10 = 0.078 MPa; k1 = 0.153 MPa; k2 = 0.109. Dispersion, κ, was set to zero for all fibre models except for the isotropic model, where it was set to 0.333. For the model without fibres, a Neo Hookean material definition was used with C10 = 0.078.

### Design of experiments (DOE)

DOE was used to identify the best combinations of structural components in the leaflet. Three structural features with four discrete levels were investigated: number of MEW layers (2, 5, 8, 10), fibre spacing (0.5, 1, 1.5, 2 mm), and silicone thickness (0.2, 0.3, 0.4, 0.5 mm). These parameters were chosen based on feasible manufacturing ranges for layers and fibre spacing, and based on bioprosthetic valve leaflet thickness (Armfield et al. 2024; Bruschi et al. 2013; Whelan et al. 2021). JMP (JMP Pro 18, NC, USA) was used to map the DOE runs; a response surface design was chosen with measured responses and targets shown in Table 1. Each response was weighted equally, except for average opening area and closed area which were weighted 2x as these are the only features that can be assessed clinically. Thirteen runs, 12 unique and 1 repeated combination, were identified to fully map this space, they are shown in Table 2.

**Table 1.** Responses investigated using DOE, with objective for each and target, if applicable.

| **Response** | **Objective** | **Target** | **Weight** |
| --- | --- | --- | --- |
| *Average stress (belly and suture edge)* | Minimise | -- | 1x |
| *Average strain (belly and suture edge)* | Minimise | -- | 1x |
| *Maximum opening area* | Maximise | Minimum: 1.7 cm<sup>2</sup> | 1x |
| *Average opening area* | Maximise | Minimum: 1.7 cm<sup>2</sup> | 2x |
| *Average closed area* | Minimise | 0 cm<sup>2</sup> | 2x |
| *Time to opening* | Minimise | -- | 1x |

**Table 2.** Embedded MEW structure combination runs as indicated by DOE.

| **Number of layers - L** | **Fibre spacing - F (mm)** | **Thickness - T (mm)** |
| --- | --- | --- |
| 2 | 0.5 | 0.5 |
| 2 | 0.5 | 0.2 |
| 2 | 1.5 | 0.3 |
| 2 | 2 | 0.5 |
| 5 | 1 | 0.4 |
| 5 | 1 | 0.4 |
| 5 | 2 | 0.2 |
| 8 | 0.5 | 0.3 |
| 8 | 1.5 | 0.5 |
| 8 | 1.5 | 0.3 |
| 10 | 0.5 | 0.5 |
| 10 | 1 | 0.2 |
| 10 | 2 | 0.4 |

### Statistical analysis

All statistical analysis related to the DOE was performed within the JMP Pro software, and p≤0.05 was considered significant.

## Results

### Fibre orientation analysis

Maximum principal stress and strain plots of the five different fibre-reinforced leaflets are shown in Figure 6. Across all the valves, there were high levels of strain and low levels of stress. Both the bioinspired and circumferential orientations resulted in valves that achieved full closure with minimal pinwheeling, meanwhile there was excessive leaflet deformation in the other three orientations. Notably, the unreinforced, matrix only leaflets reached a peak of 119.4% strain in the belly region, and the leaflets with radially aligned fibres reached 102.6% at the commissure edges. In the other three configurations, peak strains are far lower and all at the bottom suture edge, with the isotropic configuration reaching 74.6% maximum strain, circumferential reaching 69% maximum strain, and the bioinspired orientation reaching 66.4% maximum strain. Peak stresses in the leaflets occurred in the same regions. The three leaflets with highest strain experienced the lowest peak stresses (radial: 1.25 MPa; isotropic: 1.34 MPa; matrix only: 1.57 MPa) while the bioinspired (2.34 MPa) and circumferential (2.53 MPa) orientations experienced higher peak stress.

**Figure 6.**
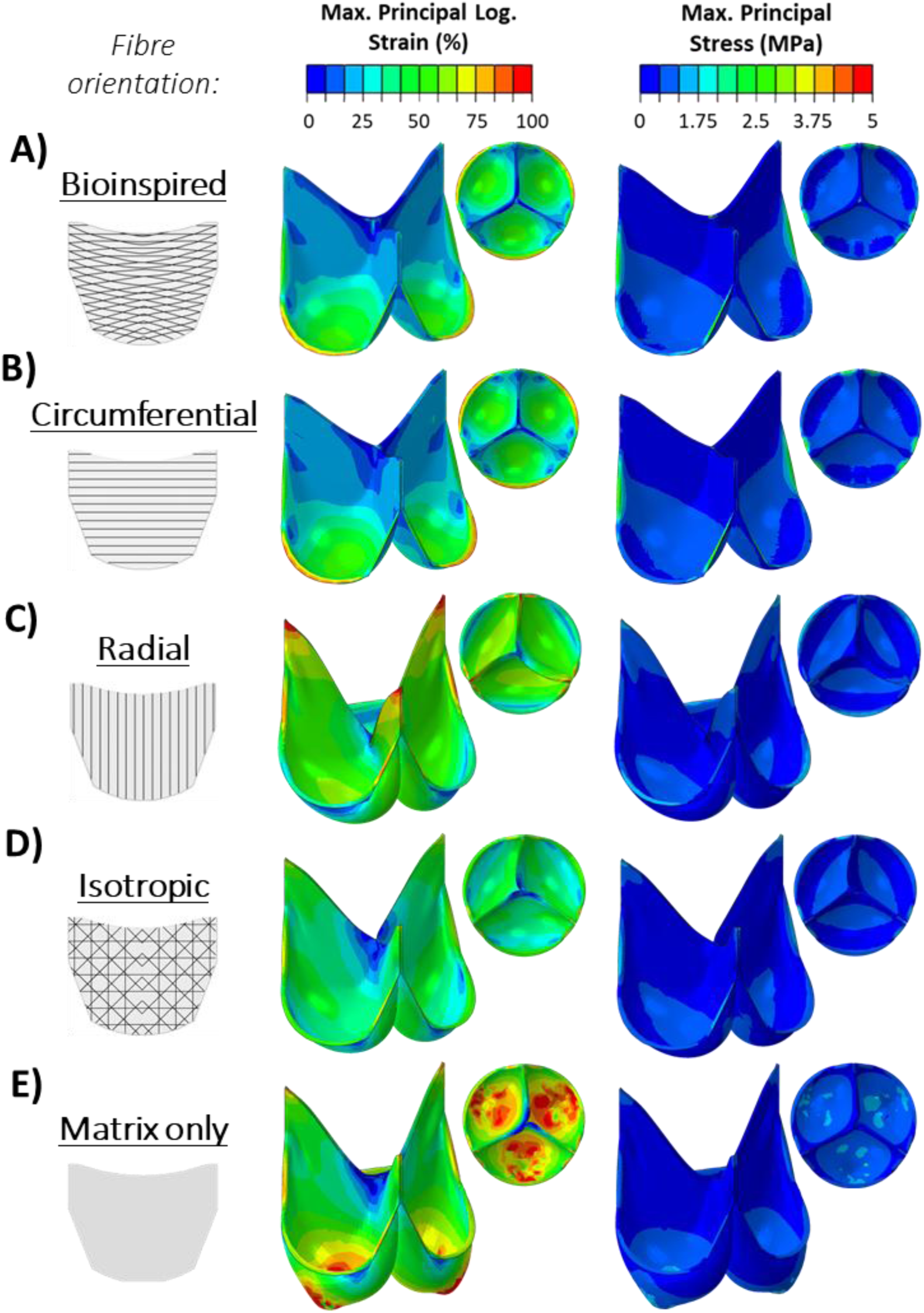
Strain and stress outputs from fibre orientation analysis showing (A) bioinspired orientation, (B) circumferential, (C) radial, (D) isotropic, and (E) matrix only with no fibre reinforcement.

Average stress and strain within the three leaflet regions is shown in Figure 7. Stress was highest in the belly region for all models, with the bioinspired, circumferential, and matrix only models experiencing the highest average stress. In the free edge, the highest average stress was seen in the radial leaflets, while in the suture region all of the leaflet configurations experienced similar levels of stress. Greater differences between the reinforcement configurations can be seen in the strain levels, with both the bioinspired and circumferential models experiencing lower strain than the other three models. In particular, the radial configuration and matrix only models experienced greater strains across all three regions.

**Figure 7.**
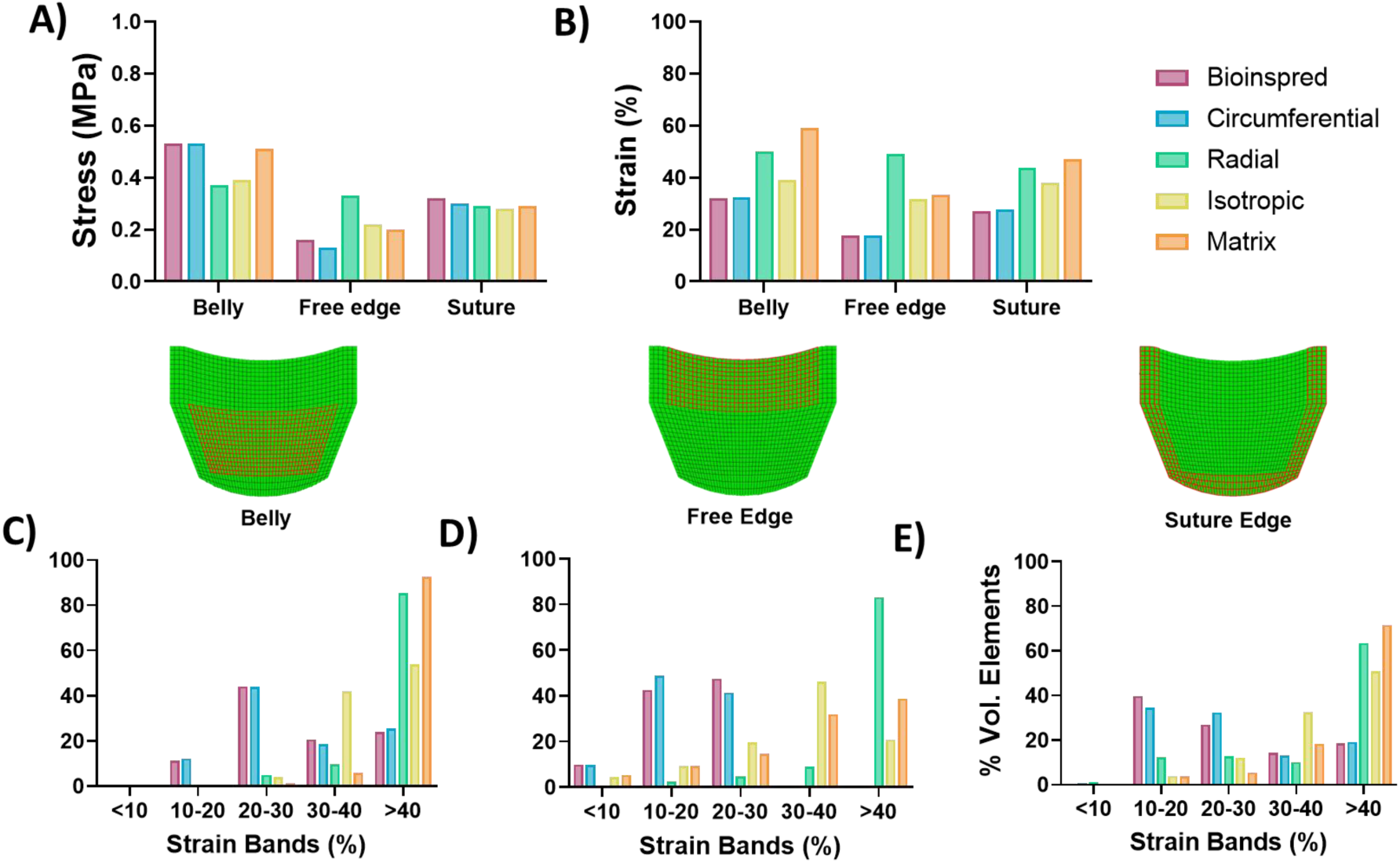
Stress and strain values extracted from models with differing fibre reinforcement structures at three different regions. (A) Average stress for each model in each region, (B) average strain for each model in each region, and volume percent of elements in different strain bands for (C) belly, (D) free edge, and (E) suture edge.

Strain in the leaflets was further investigated by organising into percent volume of elements within 10% strain bands, see Figure 7C-E. In the suture edge, the bioinspired and circumferential orientation models have a majority of the region experiencing less than 30% strain, while the other three configurations have a majority above 40%. Similar trends are seen in the free edge and the belly regions, where the bioinspired and circumferential models have more elements in the lower strain bands while the other three models have large volumes in the higher strain bands above 30 and 40%.

Opening areas throughout the cycle varied greatly between the models, with the bioinspired and circumferential configurations reaching the largest maximum OA and average OA, see Figure 8. The matrix only leaflets had the worst performance with the smallest maximum OA and average OA, with the radial configuration performing slightly better.

**Figure 8.**
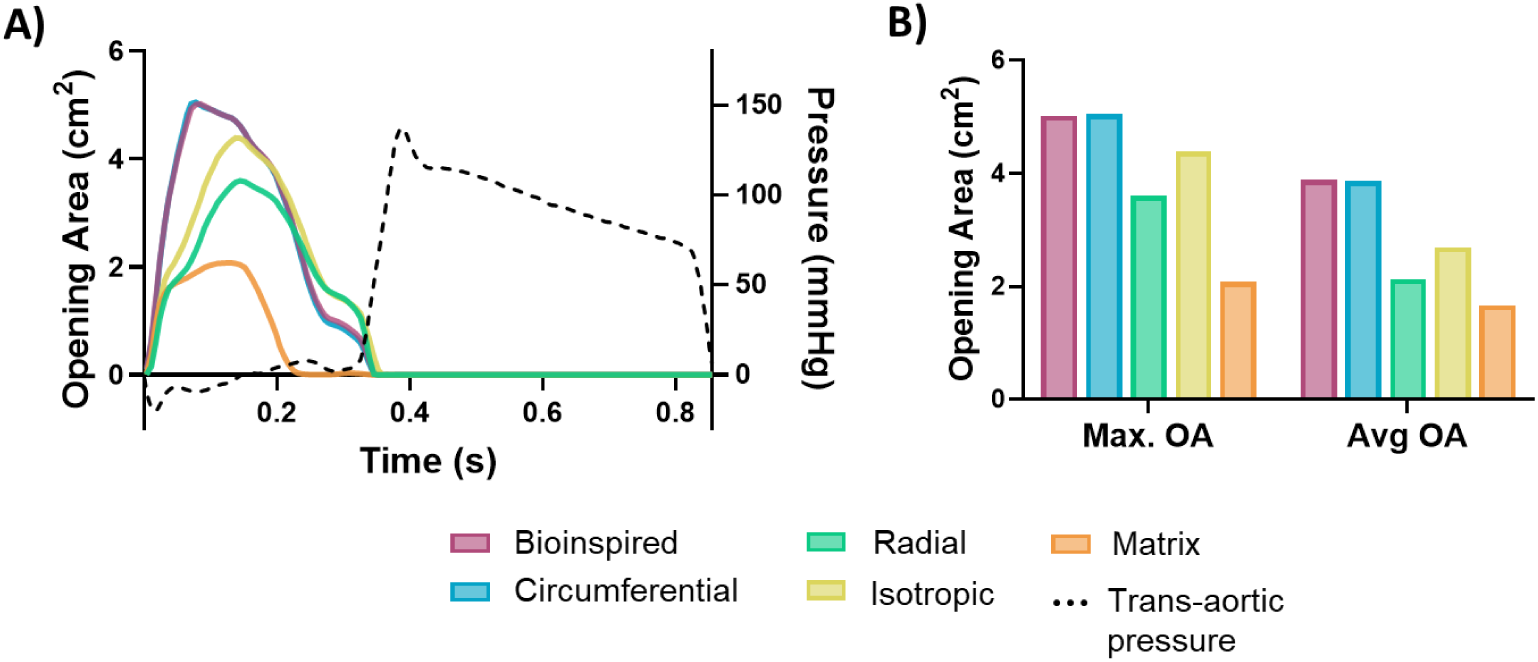
Opening area analysis for valve models with different fibre reinforcement structures. (A) Opening area for each model throughout the cycles, and (B) maximum and average opening area values for each configuration.

### Design of experiments

#### Valve models

Figure 9A shows the stress/strain response of each combination of parameters and the corresponding valve opening area throughout the cycle. A wide range of stiffnesses were simulated across the 12 unique fibre/matrix structural combinations, leading to a variation of opening area responses. Looking at the stiffest combination (10 layers, 0.5 mm fibre spacing, and 0.5 mm thickness), there were high levels of stress at the suture edge with low strain across the leaflet (see Figure 9C). Conversely, the most compliant combinations (2 layers, 2 mm fibre spacing, and 0.5 mm thickness) experienced low stress and high strain across the leaflet (see Figure 9D,E). Additionally, the more compliant material resulted in a larger maximum opening area (4.73 vs 4.20 cm^2^) and achieved full closure, where the stiffer combination did not close fully with an average closed area of 0.099 cm^2^.

**Figure 9.**
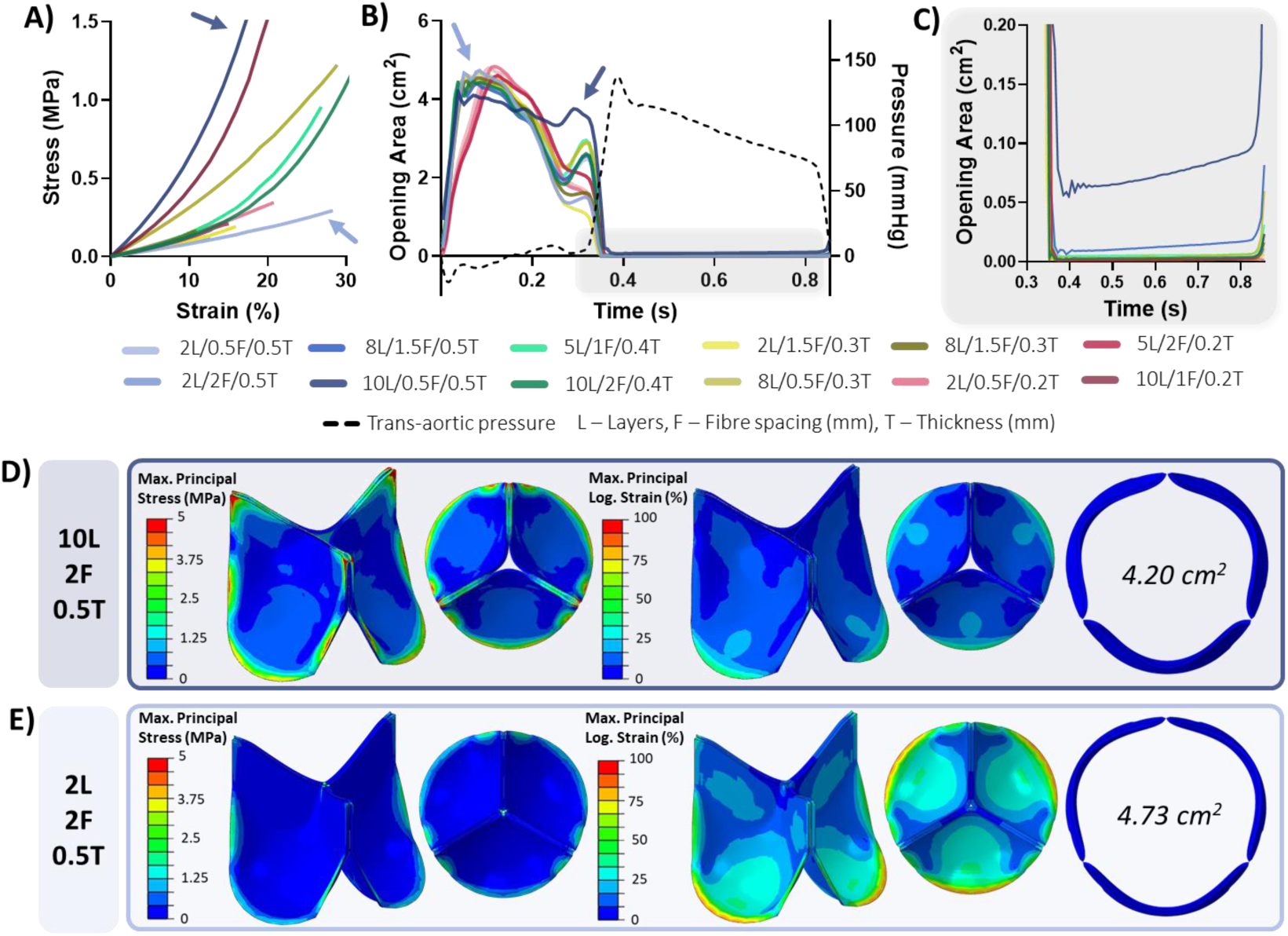
Results of all models run in DOE analysis, showing (A) stress-strain curves from uniaxial model, (B) opening areas throughout the circle, (C) detailed view of closed areas, and detailed view of valve models with (D) stiffest material response and (E) most compliant material response. Arrows indicate response of two valve examples.

#### DOE results

As seen in Table 3, the regression model was able to achieve a good fit for each response, with all R squared values reaching over 0.95. Each factor on its own had numerous significant linear relationships with the responses, with the number of layers and fibre spacing significantly impacting the leaflet stress, strain, and maximum opening area. Fibre spacing also had a significant impact on the time to reach maximum opening area, and leaflet thickness had a strong significant impact on all responses except strain at the suture edge and the average closed opening area. There are fewer significant nonlinear relationships, with all having a less strong statistical significance than the linear fit except for leaflet thickness and strain at the suture edge: this is the only nonlinear relationship with a higher P-value than the linear fit.

**Table 3.** Results of relationships between all input factors and the resulting response. Red numbers indicate a significant p-value less than 0.05, yellow numbers indicate a p-value less than 0.01. L: layer number; F: fibre spacing; T: thickness.

| Response | Fit R<sup>2</sup> | Factors |  |  |  |  |  |  |  |  |
| --- | --- | --- | --- | --- | --- | --- | --- | --- | --- | --- |
|  |  | L | L<sup>2</sup> | F | F<sup>2</sup> | T | T<sup>2</sup> | L * F | L * T | F * T |
| Suture edge stress | 0.99 | 0.0005 | ns | 0.0009 | ns | 0.0001 | 0.0322 | 0.0021 | ns | ns |
| Suture edge strain | 0.96 | 0.0175 | ns | 0.0224 | ns | ns | 0.0369 | ns | ns | ns |
| Belly stress | 0.99 | 0.0046 | 0.0323 | 0.0326 | ns | 0.0002 | 0.0085 | 0.0137 | ns | ns |
| Belly strain | 0.98 | 0.0224 | ns | 0.0099 | ns | 0.0059 | 0.0159 | ns | ns | ns |
| Max OA | 0.99 | <0.0001 | ns | 0.0002 | 0.0018 | <0.0001 | 0.0008 | 0.0016 | 0.0003 | ns |
| Avg. closed OA | 0.96 | ns | ns | ns | ns | ns | ns | 0.0407 | 0.0313 | ns |
| Avg. positive OA | 0.98 | ns | ns | ns | ns | 0.0047 | 0.0057 | ns | ns | ns |
| Time to max OA | 0.99 | ns | ns | 0.0439 | ns | 0.0016 | 0.0056 | ns | 0.0158 | ns |

Combining the factors also resulted in significant relationships in some cases, with the pairing of layer number and fibre spacing having a significant impact on stress in both regions of interest, as well as the maximum opening area and closed opening area. Number of layers and thickness together also had a significant effect on maximum opening area and closed opening area, as well as the time to reach maximum opening.

Figure 10 shows scatter plots for every factor and response pairing that had either a significant linear or nonlinear relationship. Stress in the two regions decreased with fewer layers, greater fibre spacing, and increased leaflet thickness (Figure 10A,C). A smaller fibre spacing and increased layers reduced average strain (Figure 10B, D). A significant relationship was found between maximum opening area and all three factors, but the change in opening area for every factor was very small (Figure 10E). Average opening area was increased with a larger thickness, rising by 15.1% between 0.2 and 0.5 mm (Figure 10F). Notably, all of the maximum and average opening areas recorded were well above the 1.7 cm^2^ target. Finally, the time to maximum opening area was reduced with a higher leaflet thickness and a smaller fibre spacing (Figure 10G).

**Figure 10.**
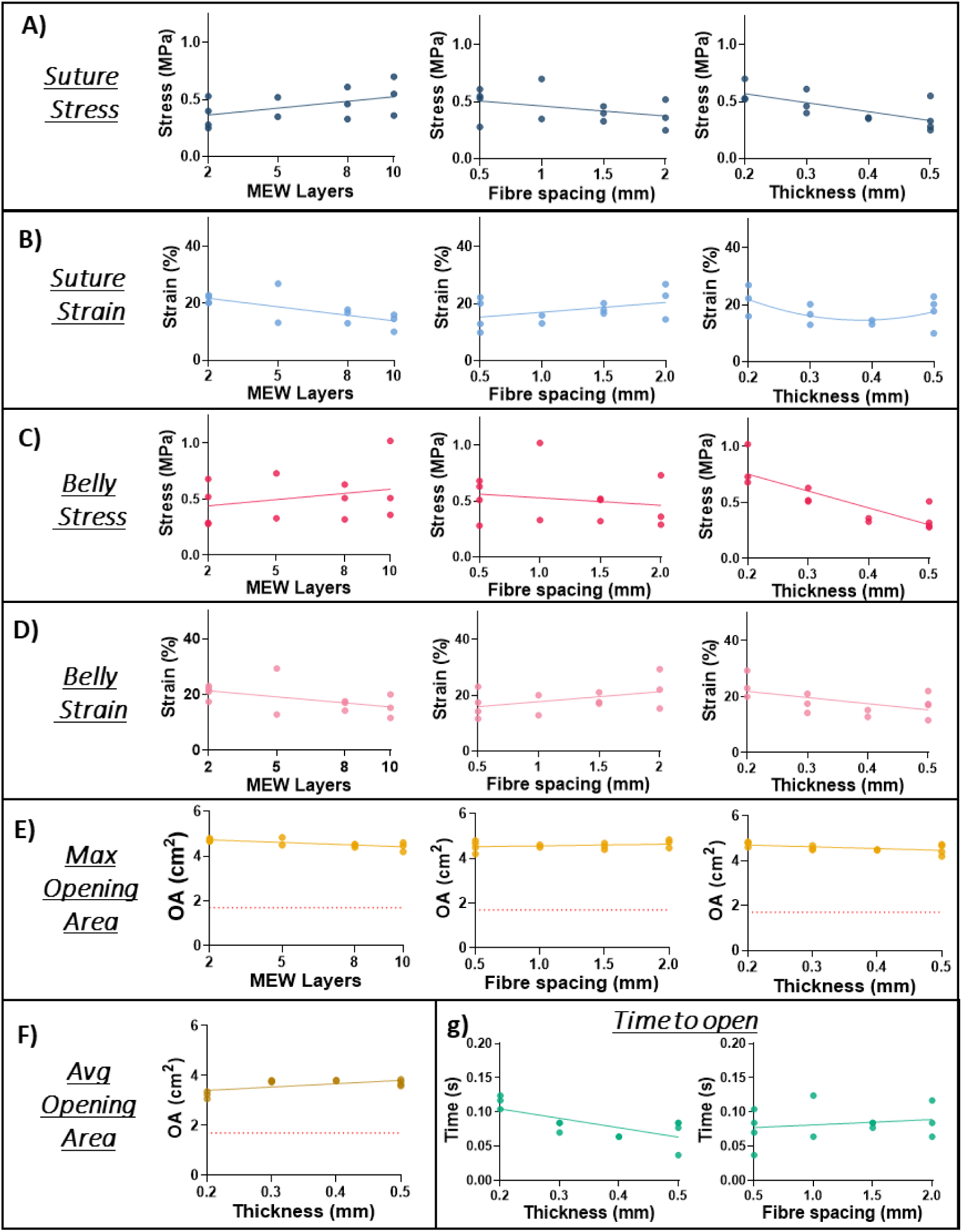
Scatter plots for every factor and response with a statistically significant relationship with an indicative line of best fit. (A) Suture stress, (B) suture strain, (C) belly stress, (D) belly strain, (E) maximum opening area, (F) average opening area, and (G) time to open.

Pairings of factors and their result on responses is shown in Figure 11. These followed the same relationships described previously, but show an extra level of complexity in the response to these two factors. In both regions, stress was increased with more layers and a smaller fibre spacing, although it was greater when the fibre spacing was 1 mm than either 0.5 or 1.5 mm (see Figure 11A,B). Maximum opening area was increased with fewer layers, a smaller thickness, and a larger fibre spacing (see Figure 11C). Time to maximum opening was reduced with a greater thickness and more layers (Figure 11D). Closed area was the only response to have a significant relationship with pairs of factors, showing a smaller area for a lower thickness, and generally fewer layers, although a large number of layers with a larger fibre spacing also improved leaflet closure (Figure 11E). It should be noted, however, that all of the valves were effectively closed, and the recorded areas were a small number of pixels.

**Figure 11.**
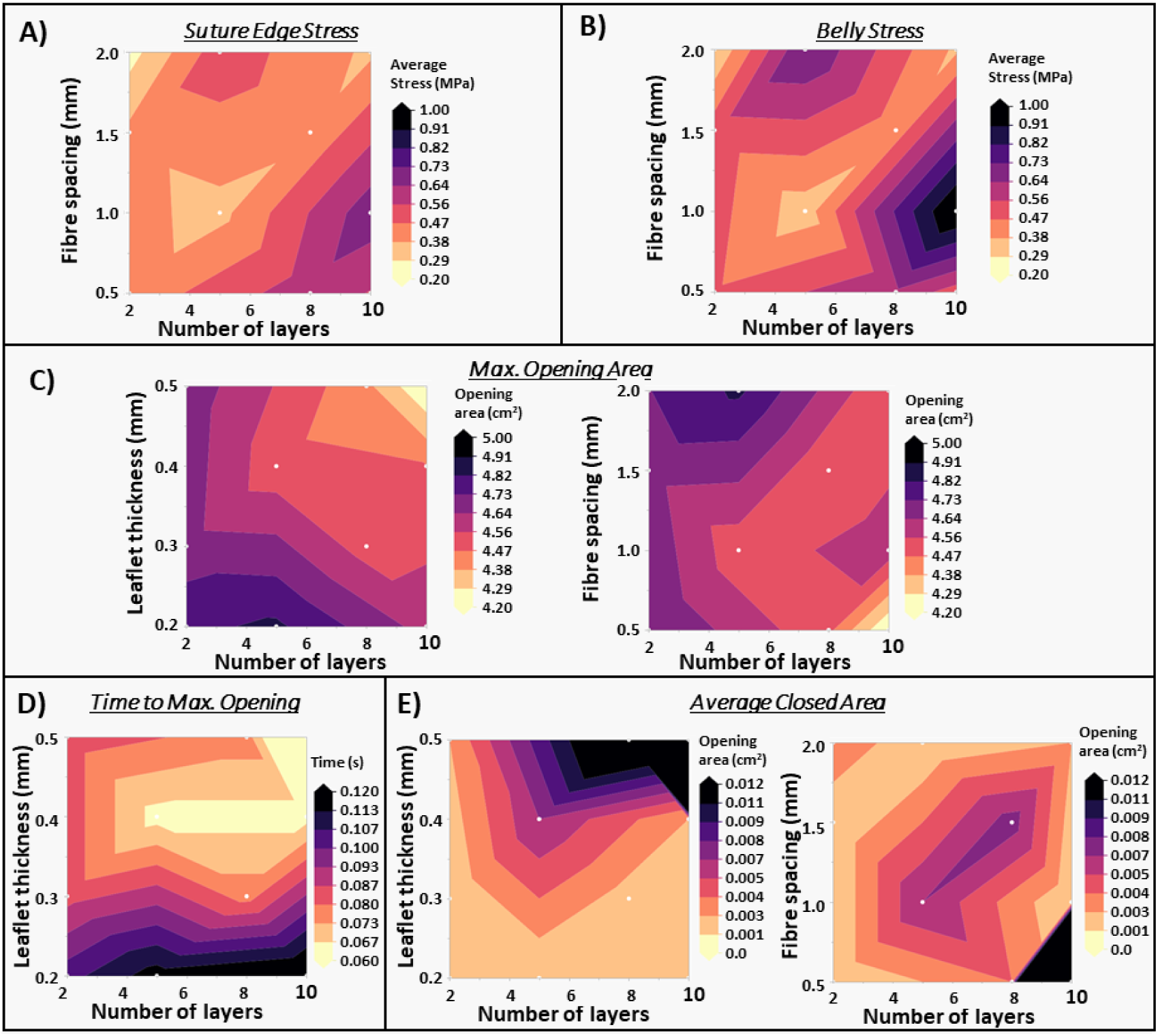
Heatmaps of all paired factors and responses with statistically significant relationships. (A) Suture edge stress, (B) belly stress, (C) maximum opening area, (D) time to maximum opening, (E) average closed area.

#### Most desirable combinations

From the results of all combinations, a “most desirable” combination of 2 layers, 0.5 mm fibre spacing, and 0.4 mm thickness was identified (named MD04, desirability: 0.78), see Figure 12A. Due to the desire to maintain a low profile device, the most desirable combinations for 0.3 mm and 0.2 mm thickness were also identified: 2 layers, 0.5 mm fibre spacing, 0.3 mm thickness (MD03, desirability: 0.65); and 5 layers, 0.5 mm fibre spacing, and 0.2 mm thickness (MD02, desirability: 0.1), see Figure 12B,C. These three combinations, not in the original DOE run, were created, run, and their results were compared to the software predictions. Average stresses in the leaflet suture edge and belly increased with decreasing thickness, while strain decreased in some regions and increased in others (see Figure 12D,E). These stress and strain measures were predicted well by the DOE, with each result falling within the predicted confidence intervals for each response. Additionally, the DOE correctly predicted an increase in stress in these regions with decreasing leaflet thickness. Strain levels in each of these regions also fall within or very close to the predicted intervals; however, MD02 experienced slightly lower average strain in these regions compared to MD04 and MD03 (MD02: 18.1% suture/20.1% belly, MD03: 21% suture, 20.9% belly, MD04: 21.5% suture/19.9% belly), where the DOE predicted the strain would be slightly higher than the other two combinations (MD02: 20.5% suture/21.74% belly, MD03: 17.22% suture/16.49% belly, MD04: 16.21% suture/14.37% belly), see Figure 12E.

**Figure 12.**
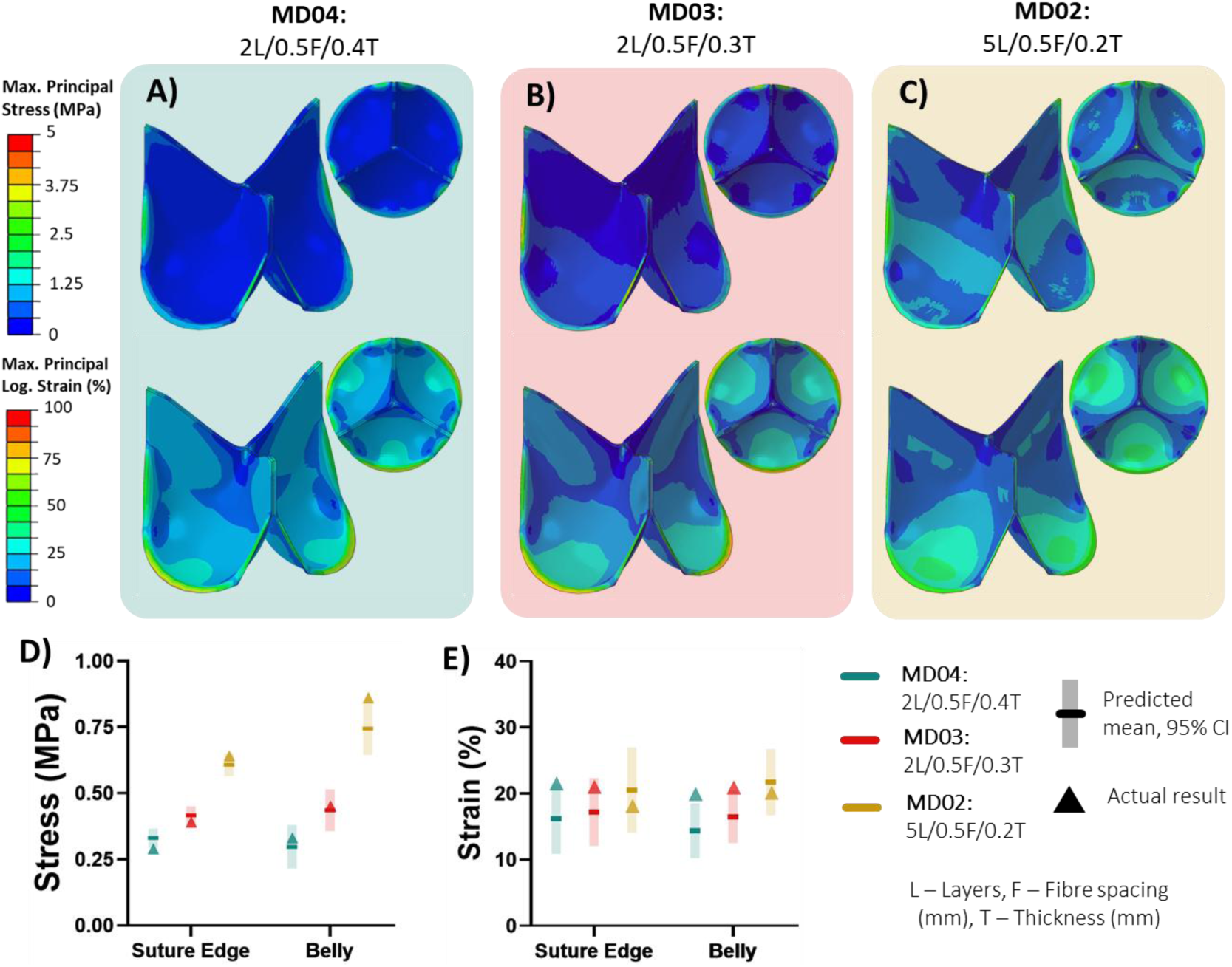
Stress and strain results of three most desirable (MD) combinations: 2 layers, 0.5 mm fibre spacing, 0.4 mm thickness (MD04); 2 layers, 0.5 mm fibre spacing, 0.3 mm thickness (MD03); and 5 layers, 0.5 mm fibre spacing, 0.2 mm thickness (MD02). Visualisations of stress and strain in leaflets for (A) MD04, (B) MD03, and (C) MD02, as well as results compared to DOE predictions for (D) stress in the suture edge and belly regions, and (E) strain in the suture edge and belly regions for all combinations.

Opening areas across the cycle for each most desirable combination are shown in Figure 13A. The maximum area reached was similar for each valve (MD04: 4.82 cm^2^, MD03: 4.81 cm^2^, MD02: 4.7cm^2^), while MD02 had a lower average opening area than the other two combinations (MD04: 4.01 cm^2^, MD03: 3.89 cm^2^, MD02: 3.67 cm^2^), see Figure 13B,C. Additionally, MD02 took slightly longer than the other two combinations to completely open (MD04: 0.07 s, MD03: 0.084 s, MD02:0.097 s), see Figure 13D. All three valves reached full closure. These behaviours were predicted well, with all of the results for maximum opening area, average opening area, average closed area, and time to maximum opening falling within the predicted intervals and very close to the mean (see Figure 13B-D).

**Figure 13.**
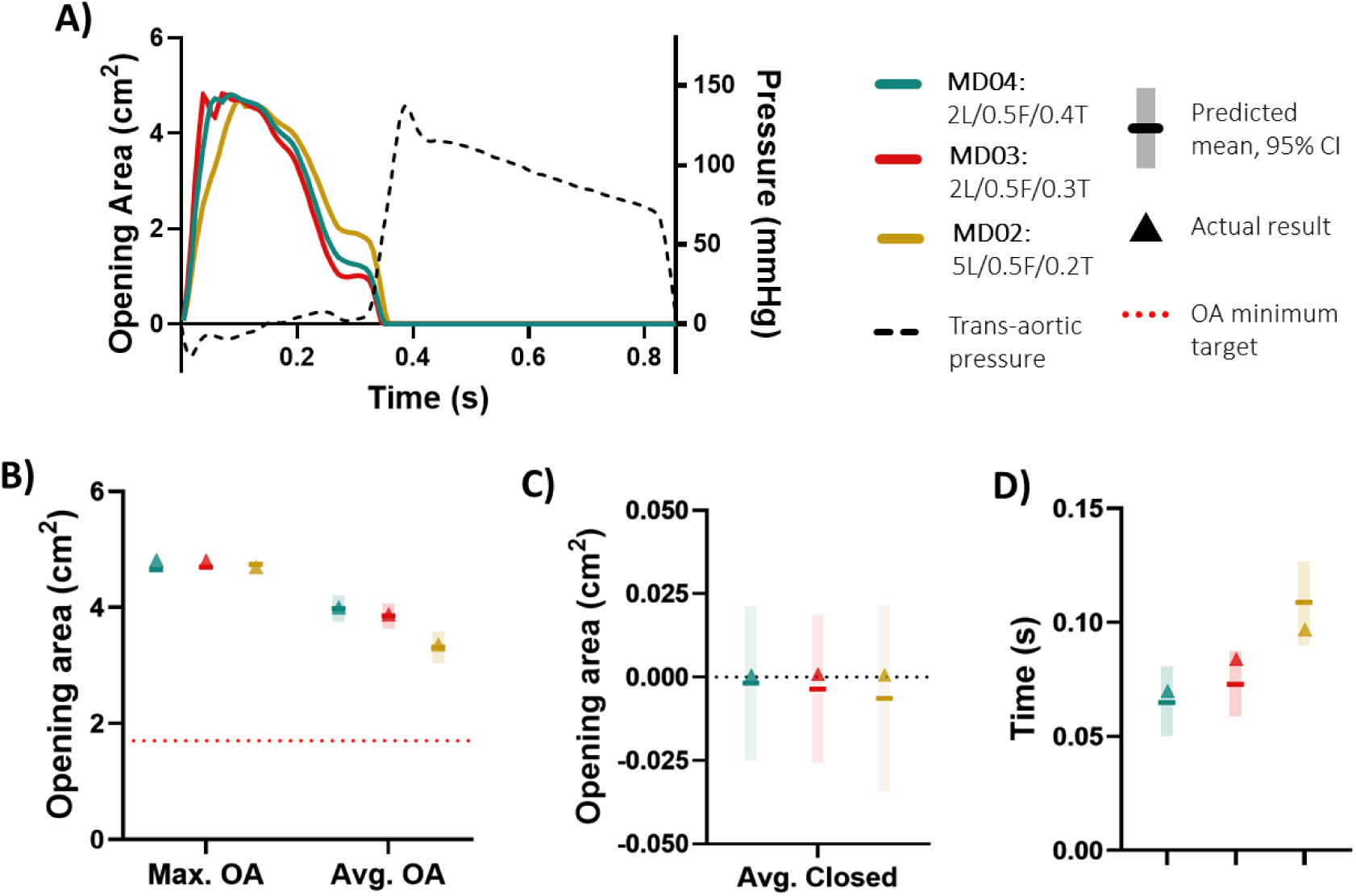
Opening area results for the three most desirable combinations, showing (A) opening areas versus pressure for a cycle, (B) maximum and average opening area results versus predictions, (C) average closed areas versus predictions, and (D) time to maximum opening versus predictions.

## Discussion

This work presents the creation of a finite element and design of experiments (FE-DOE) based framework to aid the design of future bioinspired polymeric aortic heart valves. Using a fibre-reinforcement structure inspired by MEW, we developed a methodology to obtain the mechanical behaviour of PEEK fibre-reinforced PDMS and simulate this material in a trileaflet valve model. After showing the positive impact of implementing a bioinspired fibre structure in this leaflet geometry, this methodology was used alongside DOE to determine the best structural combination of a PEEK-like fibre structure manufactured using MEW and embedded in PDMS.

The first step was to determine the stress/strain response of a PEEK fibre-reinforced PDMS in silico using a discrete fibre-reinforced dogbone model simulating a uniaxial tensile test. The response of this composite material was used to calibrate the relevant parameters of the GOH fibre-reinforced model which significantly reduced computational complexity and simulation time. Pressure was applied to the aortic and ventricular surfaces according to measurements taken from pulse duplicator testing of size large ACURATE neo2 valves. The pressure curves show some minor differences from the commonly referred Wiggers Diagram curve which is most likely due to system setup and the resolution of the pressure measurements, but they align with other pressure-time curves given in literature on bioprosthetic valves (Armfield et al. 2024; Wu et al. 2019).

We showed the flexibility of this model and the impact of fibre orientations on material response and valve performance by simulating leaflets with varying reinforcement configurations. These results clearly showed the impact of different reinforcement structures on material strain and valve performance While the bioinspired and circumferential orientation models performed similarly, the bioinspired orientation induced lower peak strain and stress at the suture edge which could help extend material lifetimes. Additionally, the impact of changing leaflet fibre orientations shows the positive impact of customised, local fibre reinforcement. These insights are corroborated by a study performed on similar leaflet geometry for porcine pericardium by Armfield, et al., where the authors showed higher leaflet displacement for radially aligned fibres compared to circumferentially aligned fibres (Armfield et al. 2024). Also, Serrani et al. found a reduction in overall and maximum leaflet strain with an optimised fibre reinforcement structure compared to an isotropic orientation (Serrani et al. 2016).

For the analysis of these valve models, a robust and repeatable assessment framework was established to provide broad insights into leaflet performance and enable future design choices..

Using DOE we were able to determine the impact of fibre spacing, number of layers, and overall thickness on leaflet and valve performance A variety of responses were investigated, including both material behaviour and clinically-relevant performance indicators of opening area.

All of the models achieved good opening areas, far surpassing the 1.7 cm^2^ minimum for this size valve in both the maximum OA and average OA (ISO5840-3). Additionally, all but the stiffest material combinations reached full valve closure. While the materials in each of these models may differ in their response, these opening areas suggest that these stiffness ranges are good candidates to enable effective aortic valves.

Greater differences in the leaflet structural combinations were seen in the stress and strain results, particularly in the belly region. In order to ensure a durable and long lasting leaflet, the stress and strain in the material should be minimised. In addition, the models differed in their time to reach the maximum opening area, particularly for the smaller thickness leaflets, suggesting poorer valve efficiency. Using the DOE results we can see that leaflet thickness had a significant impact on these responses, with a larger thickness leading to reduced stress, strain, and time to maximum opening area. Fibre spacing and number of layers also had a significant impact on the leaflet stress and strain, but in opposing ways: fewer layers and a larger spacing increases strain while decreasing stress. A larger fibre spacing will also increase time to maximum valve opening. Notably, there are no independent factors that impact valve performance and material response more than the others, all three elicit significant impacts on many of the responses. Meanwhile, almost all of the valves fully close, therefore none of the factors alone significantly impacted this. While two combinations of factors (layer number and fibre spacing; layer number and thickness) did result in a significant correlation to closed area, the actual values of the closure area are so small that this means very little.

The overall complexity in how each of the factors contributed to the responses argues for using this DOE approach as it is able to balance the interactions between each of these factors with the desired outcome for each factor (i.e. maximise or minimise) to provide a “most desirable” combination. We were able to identify the most desirable structural combinations for the leaflets based on the input criteria outlined in Table 1. The most desirable combination identified was 0.4 mm thick, which is thicker than porcine pericardium used in many self-expanding TAVR valves (Armfield et al. 2024; Bruschi et al. 2013).Due to the clinical need for a thinner leaflet to maintain a low profile device when crimped, we also identified the best combinations at the two smaller thicknesses: 0.3 and 0.2 mm. These combinations were created, calibrated, and run in the dynamic valve model, and their results showed a close agreement with the DOE predictions. All of the responses fell within the 95% predicted confidence interval bands, and close to the mean, except for belly strain in the MD04 and MD03 models whose actual values were slightly above the maximum predicted. Additionally, it was predicted that MD02 would experience higher strains than MD04 and MD03 which was not the case. Outside of this, the rest of the responses were predicted very well.MD03 achieved similar but overall slightly worse results to MD04 across all categories, which suggests that in order to maintain a low device profile with thinner leaflets there is a necessary trade-off with the mechanical response of the material. From these results and taking into account the industrial need for thinner leaflets, the best combination from this dataset is likely MD03, as it is the thinnest option with the least compromise on material response.

This work does contain limitations. First, a frame was not included in the model. It is known that the frame in the ACURATE neo2™ displaces under leaflet loading, which would impact the stress and strain in the leaflets(Armfield et al. 2024), and potentially the opening and closing behaviour. Fluid flow was not included either, which is likely to impact the same behaviours. While future work should investigate the impact of frame deflection and fluid flow inclusion, it is unlikely to change the DOE outcome as all the leaflet combinations are compared relative to each other in the same conditions. Second, the fibres were modelled as a linear elastic PEEK-like material with no yield included. While PEEK is a stiff polymer, it also has a low yield strain (Shrestha et al. 2016). Future work should look at the impact of this, and investigate using PCL as this is the most used and accessible material used for MEW. Additionally, the GOH model is used to describe the fibres in the leaflet material rather than including discrete fibres. This could impact the material behaviour, especially with combinations containing larger pore sizes as it is not currently able to replicate local stiffening in one area and an unreinforced region elsewhere. Using this material description greatly reduces model complexity and makes the model running times feasible. Again, the overall affect across the leaflet and valve is likely to be maintained despite this simplification.

On the DOE itself, some factor combinations would not be physically possible to manufacture. For example, while a 10-layered structure made of 20 µm-diameter fibres could theoretically be embedded within 0.2 mm of silicone, this would not be possible on the bench using MEW due to the high peaks formed when fibres overlap (Cao et al. 2021; Lamb et al. 2024), resulting in a total thickness in these regions of greater than 0.2 mm. This would also pose manufacturing challenges for structures with 8 layers at the same total leaflet thickness.

## Conclusion

Overall, a FEM-DOE framework has been developed which offers a powerful and efficient methodology for identifying the best combination of MEW-based structural features and leaflet thickness for the ranges investigated and materials used. By utilising a fully in silico approach, it removed the need for manufacturing leaflets, assembling valves, and testing these devices, significantly reducing time spent identifying the best combination. While these results only hold for the materials investigated in this study, the same methodology can be applied to different materials to identify the best structural combinations for a variety of material combinations. Ultimately, this is a highly valuable tool that can allow researchers the ability to probe potential material combinations to identify the best structures for in vitro testing. Additionally, its versatility allows the investigation of a variety of factors. While layers, fibre spacing, and thickness were investigated here, other factors can be explored such as fibre diameter, matrix stiffness, and even MEW processing parameters. The value of linking FE methods with DOE allows for the structured exploration of numerous variables, while minimising time spent manufacturing and testing the chosen combinations.

## Acknowledgements

This project is co-funded by Taighde Éireann/Research Ireland and Boston Scientific Corporation (EBPPG/2020/200).

