## Supplemental material for "FE-DOE: Finite element informed design of experiments to optimise bioinspired melt electrowritten (MEW) polymeric heart valves"

**Supplementary Material**

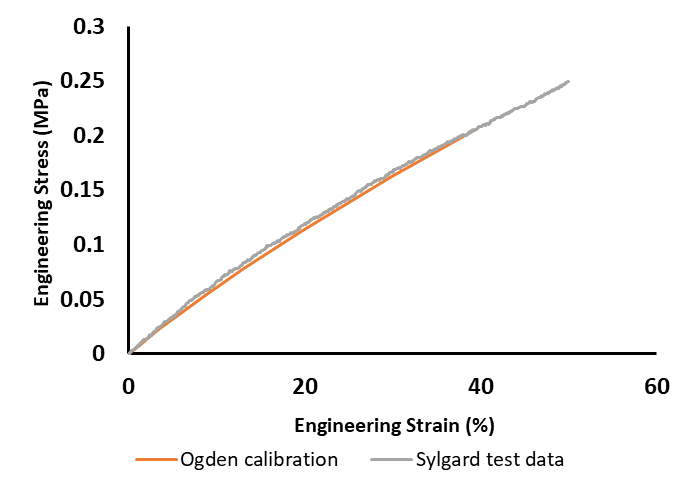

**Figure S1** Comparison of PDMS mechanical response to a calibrated first-order Ogden model (µ1= 0.204 MPa, α1= 2.504, and D=0).

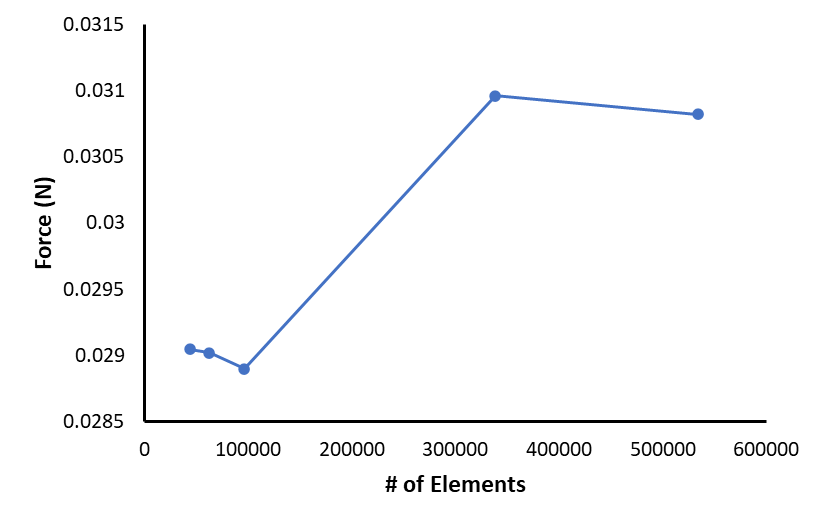

**Figure S2** Mesh sensitivity analysis of dogbone model.

Reaction force at 20% strain for a variety of element numbers. Convergence was achieved by 35,000 elements which corresponded to a seed size of 0.05 mm for the fibres and 0.2 mm for the matrix.

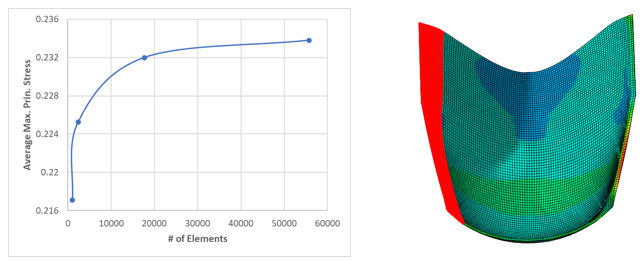

**Figure S3** Mesh size sensitivity analysis.

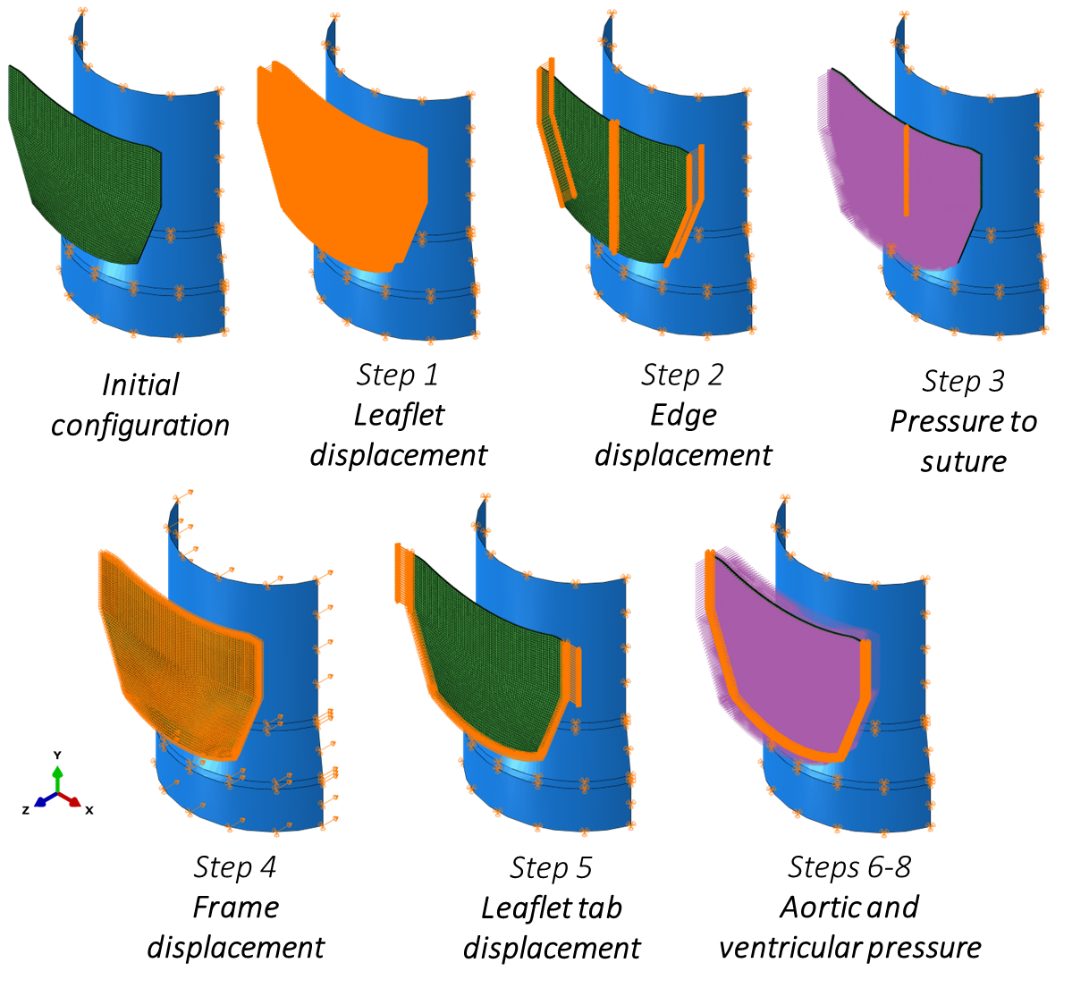
Average maximum principal stress along the suture edge was analysed for increasing mesh densities and convergence was achieve at around 20,000 elements.

**Figure S4** Detailed images of leaflet and frame boundary conditions

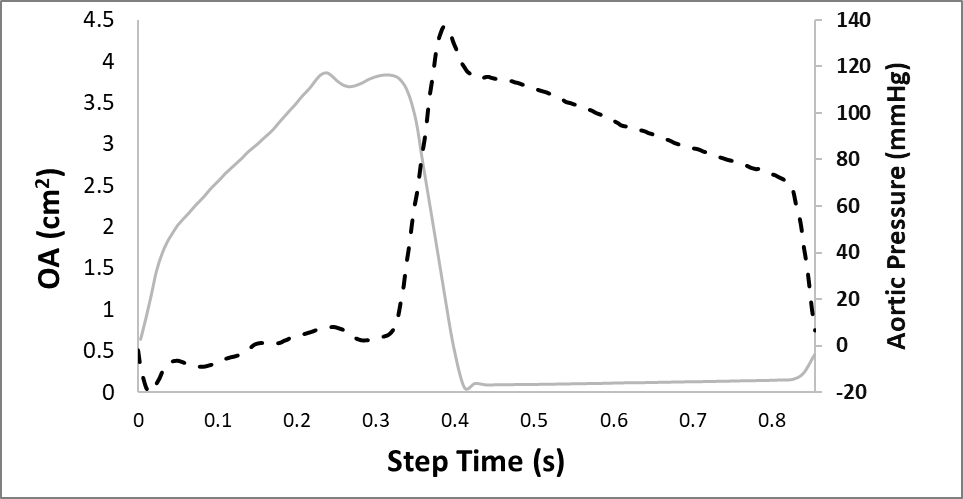

**Figure S5** Opening area of porcine pericardium model compared to resultant aortic pressure.

Material properties of highly-dispersed porcine pericardium were applied to the trileaflet valve model and the average OA was compared to expected EOA ranges provided by Boston Scientific to validate the comparability of these outputs. It was found that the valve model accurately predicted measured EOA valves, with the average OA reaching 2.14 cm^2^ compared to the expected range of 2.0-3.0 cm^2^.
